# Cancer cell hyper-proliferation disrupts the topological invariance of epithelial monolayers

**DOI:** 10.1101/2021.08.12.455991

**Authors:** Daria S. Roshal, Marianne Martin, Kirill Fedorenko, Virginie Molle, Stephen Baghdiguian, Sergei B. Rochal

## Abstract

Although the polygonal shape of epithelial cells has drawn the attention of scientists for several centuries, only recently, it has been demonstrated that distributions of polygon types (DOPTs) are similar in proliferative epithelia of many different plant and animal species. In this study we show that hyper-proliferation of cancer cells disrupts this universality paradigm and results in random epithelial structures. Examining non-synchronized and synchronized HeLa cervix cells, we suppose that the cell size spread is the single parameter controlling the DOPT in these monolayers. We test this hypothesis by considering morphologically similar random polygonal packings. By analyzing the differences between tumoral and non-tumoral epithelial monolayers, we uncover that the latter have more ordered structures and argue that the relaxation of mechanical stresses associated with cell division induces more effective ordering in the epithelia with lower proliferation rates. The proposed theory also explains the specific highly ordered structures of some post-mitotic unconventional epithelia.

## Introduction

Symmetry and topology determine the structure and laws of motion for relatively simple abiotic systems studied by physics and chemistry. In living systems, gene expression is usually considered as the fundamental mechanism controlling development and homeostasis^**1,2**^. Nevertheless, the polygonal (prismatic in 3D) shape of cells, and their highly ordered packing in epithelia, clearly demonstrate that the organization of these cell monolayers directly follows basic physical and topological rules^**3-5**^. Epithelial growth is achieved by intercalary cell divisions within a constrained volume^**6**^. Dividing cells change their mediolateral neighbors, thus maintaining the apico-basal architecture and tightness of the intact layer^**7**^, which retains both robustness and plasticity. The most striking phenomenon is the so-called topological invariance observed in almost all proliferative epithelia within phylogenetically distant organisms harboring different global architectures. In fact, during their formation, different epithelial structures converge to polygonal packings with very similar DOPTs^**8-12**^. Therefore, in very different epithelia the probabilities to observe cells with the same number of sides are approximately equal. This topological invariance is closely associated with the physiological invariance of epithelium. In all eumetazoans, epithelial structures form a selective paracellular barrier, which controls fluxes of nutrients, regulates ion and water movements, and limits host contact with antigens and microbes. This invariant fundamental function is achieved by the maintenance of the epithelial tightness throughout various morphogenetic processes, like embryonic development, organogenesis, or continuous cell renewal^**13-15**^.

Among previously published studies that identified a variety of biological and physical mechanisms underlying the universal properties of epithelia, two articles are of particular interest. Back in 1928, the DOPT in the proliferative epithelium of cucumber was first investigated^**8**^: out of 1000 epithelial cells studied, 474 were hexagonal, 251 were pentagonal, and 224 were heptagonal. Later, in an excellent article^**11**^, the hypothesis of topological invariance was formulated. It was established that the similar distributions of polygons are observed in epithelia of several more animal species, and a Markov-type theory was proposed to explain such invariance. This theory assumed that the junction that appears during cell division is always located in such a manner that it forms two additional polygonal vertices, which are necessarily on non-adjacent sides of the original polygon. It was also supposed that the probability of the junction formation is independent of the ratio between the parts in which the cell is divided. The study^**11**^ was exploited in several biophysical models of proliferative epithelia^**16-20**^. However, the theoretical part of ref.^**11**^ has several shortcomings. It cannot explain the existence of 4-valent cells, for which the observed fraction varies from 2 to 3%^**8,11**^. The supposed equiprobability of cell division into substantially different parts is not justified. The theory considers neither the possibility of cellular motility nor mechanical interactions between cells, which are crucial for the the epithelium properties^**19,21,22**^. Moreover, the cell-to-cell interactions also strongly influence the initial steps of epithelial tumorigenesis^**23**^.

Epithelial tumors (carcinomas) may remain circumscribed by the surrounding epithelium, in which case they are considered benign and can be completely and easily excised by a surgeon. However, when the tumor becomes capable of disturbing the environment topology, it acquires the ability to cross basement membranes, colonize surrounding tissues, and then turns out to be much more difficult to excise and control. Since the experimental data^**8-12**^ show that topological invariance is critical for the organization and morphogenesis of healthy epithelial tissues, an analysis of topology modification during epithelial tumorigenesis can provide a lot of information on the involvement of architectural perturbations in the very early stages of tumor transformation. Until now, the use of topological models for description and analysis of tumor structures has been limited to the utilization of tumor sections providing 2D information about 3D tissues^**24**^. Such models, however, cannot answer the principal question: what is the topological difference between tumoral and normal epithelium? Our new interdisciplinary approach applied to *in vitro* models of tumoral and non-tumoral epithelial cells of the same origin (cervix) allowed to decipher this difference and identify the physical mechanism that maintains the topological invariance of normal epithelia and ceases to operate in the tumor monolayers.

The first objective of our study is to test whether the cancer epithelium retains the topological invariance. For this, we investigate the structural characteristics of non-synchronized (conventional) and synchronized monolayers obtained from HeLa epithelial cancer cells. The latter monolayers are of particular interest, since most of the cells constituting this model epithelium belong to the second generation, and the Markov-type theory used in ref.^**11**^ goes beyond the scope of its applicability. Moreover, despite numerous previously published data^**25**^, a putative influence of the cell cycle duration and cell synchronization on the shape and topology of epithelia remains unclear^**13**^.

Having discovered that cancer epithelia are able to break the topological invariance, in order to rationalize the observed structures, we propose a new geometrical model based on a random (equiprobable) disposition of cells with polygonal shape and different sizes over the epithelium. The model generates polygonal packings that are very similar to the structures observed in non-synchronized and synchronized HeLa monolayers. Testing and applying our approach to several normal proliferative epithelia, we reveal the topological difference between tumoral and non-tumoral epithelia and propose the physical mechanism underlying this difference. As we demonstrate, in the epithelia with lower proliferation rates the relaxation of mechanical stresses associated with cell division and cell growth results in more ordered structures and maintains the topological invariance. Our theoretical approach allows to better understand the morphological stability of epithelia; in addition, it can also be used to understand various aspects^**26**^ of epithelial tumor formation.

## Results

### Structural characterization of cancer cell monolayers

Before carrying out the structural characterization of the epithelial monolayers that we obtained, it is important to discuss some general properties of DOPT. Note that the cell polygons in the epithelium are convex and form a tessellation of the monolayer surface. The tessellation is similar to the Voronoi one^**27**^, which, in turn, is dual to the specific triangulation of the surface by acute-angled triangles. In this case, for an infinite arbitrary flat monolayer, the average number of nearest neighbors equals 6. Indeed, if the Gaussian curvature is absent, then the equality *Δ*=0 takes place, where

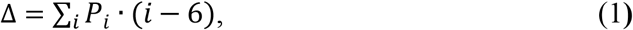

*P*_*i*_ = *N*_*i*_ / Σ_*j*_ *N*_*j*_ is the concentration of cells with *i* nearest neighbors and *N*_*j*_ is the total number of cells, with *j* nearest neighbors. Recall that the quantity *Q*_*i*_ = Σ_*j*_ *N*_*i*_ · (*i* − 6) proportional to *Δ*, is called the topological charge^**28-30**^. For a triangulation of the sphere, *Q* = 12, for a torus, as well as for an infinite plane, *Q* = 0.

In fact, the non-local equality *Δ*=0 or the equivalent statement about the average number of neighbors means that the DOPT must be balanced: the number of cellular *n*-gons, with *n*<6, must balance the number of *n*-gons with *n*>6. In this equilibrium, the weights of 5- and 7-gons are equal to one, the weights of 4- and 8-gons are twice as large, etc. Eq. (1) can be used to estimate the error in the experimental determination of DOPT. For a finite monolayer, or when averaging over several samples, the obtained value of *Δ* can deviate from zero, characterizing the error of the experimental method for calculating the DOPT. In particular, due to the relatively large number of cells considered in *Cucumis* epithelium^**8**^, the error (1) for this case is *∼*0.004, which is almost 10 times lower than for the data in ref.^**11**^. Nevertheless, in the geometrically correct model of cell division^**11**^, with an increase in the number of successive divisions, *Δ* tends to 0, reaching *Δ*≈0.001 at 10^th^ division. Since both the left and right sides of the distribution contribute to *Δ*, it is reasonable to estimate the maximum error in determining the probabilities *P*_*i*_ in the DOPT as *Δ*/2. Also, the condition *Δ*=0 severely restricts the possible forms of the DOPT. In real epithelia, probabilities *P*_*i*_, where *i*>7 or *i*<5, are small. The critical probability *P*_6_ is, and other probabilities follow it conserving the condition *Δ*=0. For example, if in a hypothetical epithelium consisting only of 5-,6- and 7-valent cells, the percentage of hexagonal cells is *P*_*6*_ then concentrations of 5- and 7-valent cells should be equal to (1-*P*_*6*_)/2.

HeLa cells are human malignant epithelial cells derived from an epidermoid carcinoma of the cervix. The growth of HeLa cell monolayers and synchronization procedure are described in Methods, see also Supplementary Fig. 2. In synchronized monolayers, most of the cells belong to the second generation with similar time elapsed after the division process. The characterization of the HeLa and HeLa synchronized epithelia was carried out using 9 and 7 assembled images obtained by the juxtaposition of contiguous microscope fields. The studied epithelial areas contained from 404 to 933 cells. The first line of Fig. 1 shows typical non-synchronized and synchronized HeLa epithelial cells (respectively samples HeLa9 and HeLa_syn_5 in Table I). Due to visualization specificities (see Methods), the cell nuclei in micrographs are clearly visible, while the cell boundaries are often poorly distinguishable. Therefore, in order to determine the number of nearest neighbors and obtain additional structural data, we used the Voronoi tessellation (see Methods), with the nodes located at the centers of the nuclei. Cells for which it was impossible to unambiguously determine the boundaries were not considered. Therefore, we took into account only those cells that fall inside the smaller rectangular area inside the assembled image (see Methods).

**Table I.**
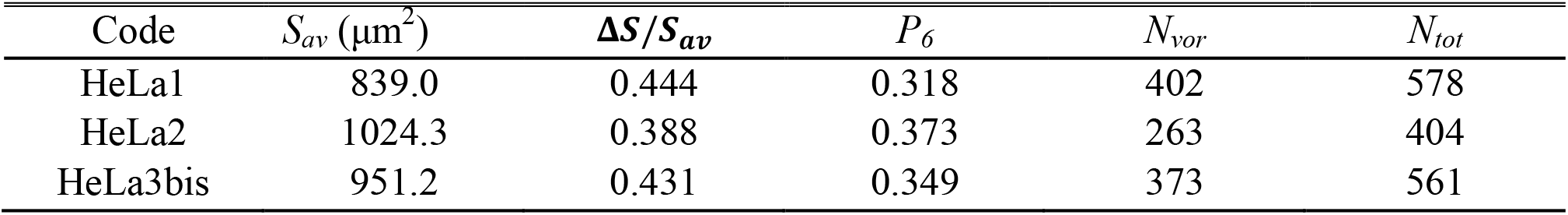
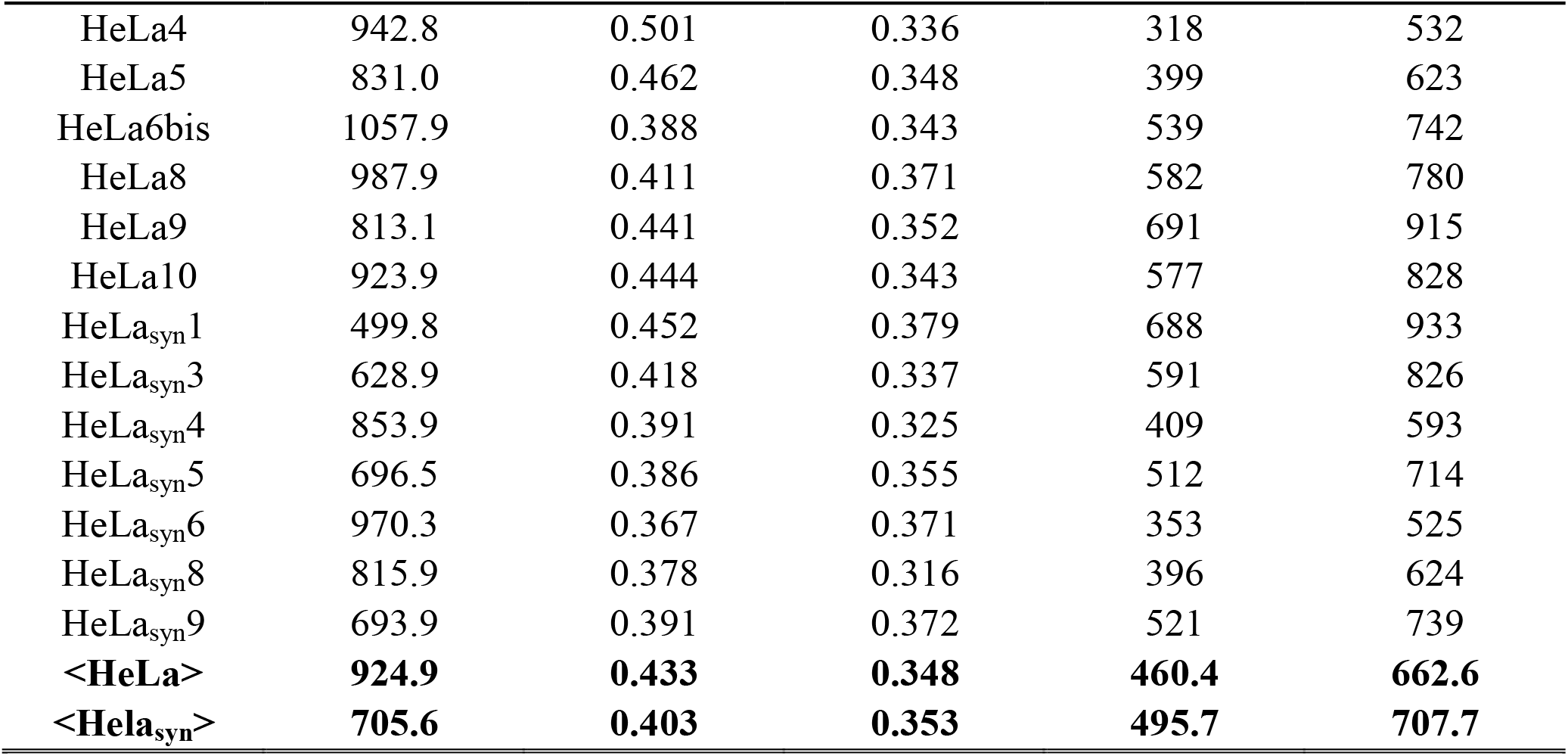
Characterization of the studied samples. The columns contain: sample name, average area of Voronoi cells *S*_*av*,_ dimensionless effective spread of areas Δ*S*/*S*_*av*_, probability *P*_*6*_, number of identified Voronoi cells *N*_*vor*_, total number of cells *N*_*tot*_. The last two lines in bold correspond to the averaged structural data for HeLa and HeLa synchronized samples.

**Fig. 1.**
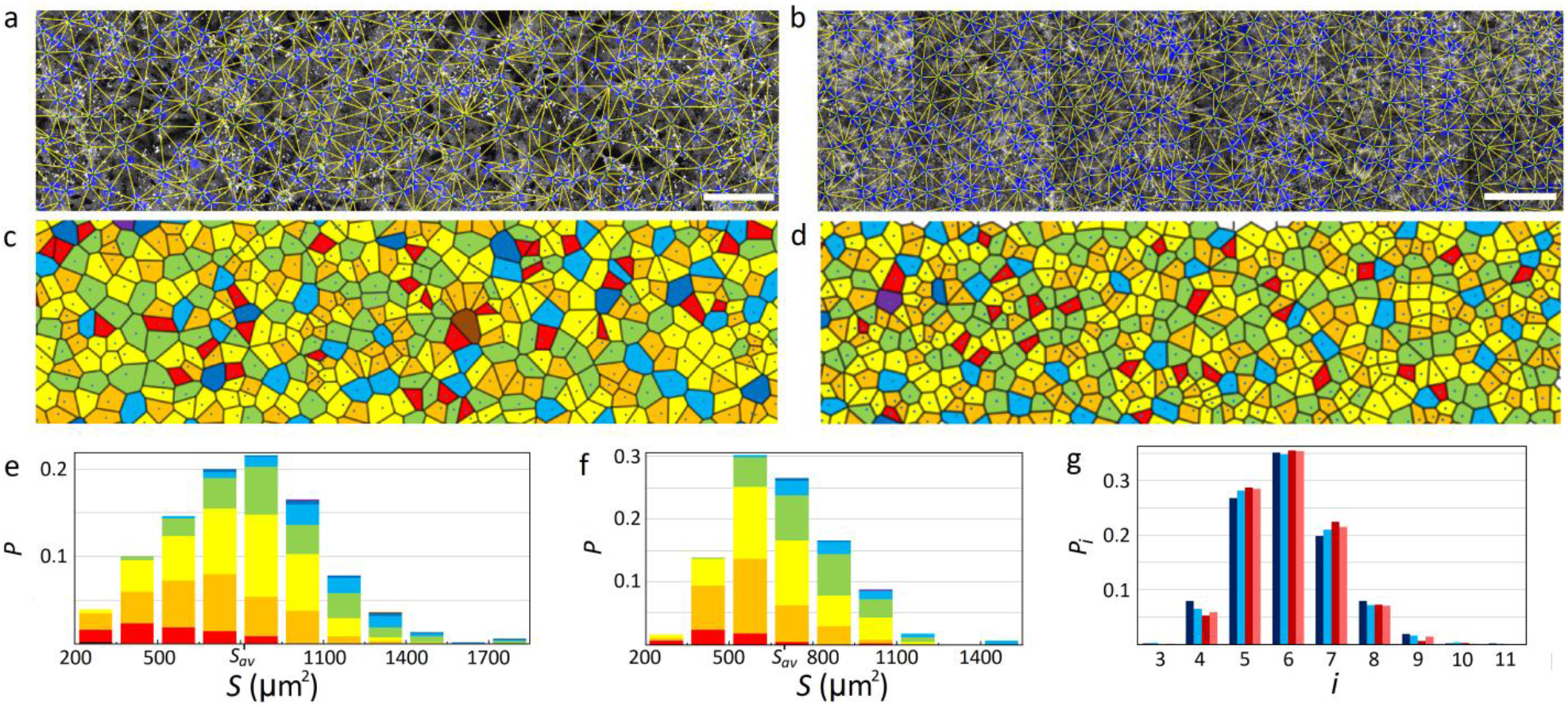
**Structural characterization of HeLa cell monolayers;** for details regarding their growth see Methods and Supplementary Fig. 1, 2. **a** Non-synchronized and **b** synchronized cellular structures. White scale bars are 100 μm. The triangulation with nodes in the centers of cell nuclei is imposed on the monolayers. **c** and **d** Voronoi tessellations for the monolayers **a** and **b**, respectively. Polygons with 3-11 sides are colored with black, red, orange, yellow, green, light blue, dark blue, purple and brown colors, respectively. Histograms **e** and **f** show a probability *P* to observe the certain ranges of cell areas *S* in the samples **a** and **b**. Contributions from different types of polygons are shown with the same colors as in **a-b. g** DOPTs for the samples **a-b** and averaged data. Dark blue, blue, red, and pink colors correspond to HeLa9, <HeLa>, HeLa_syn_5, and <HeLa_syn_> lines from Table I.

Second line of Fig. 1 shows two Voronoi tessellations for the considered cellular structures. The areas of the epithelial cells were calculated as the areas of the cells of the Voronoi tessellations. Figures 1e-f contain the histograms of cell area distributions for the considered samples. Finally, the last panel in Fig. 1 shows the DOPTs for the structures a-b, as well as the averaged distributions for HeLa non-synchronized and HeLa synchronized monolayers. In all cases, the averaged topological error *Δ* is less than 0.01.

It is important to note that all histograms of cell area distributions (including the samples not shown in Fig. 1) are wide and correspond to the Gauss type, which can be explained by the spread of the cell sizes before the mitosis, and the possibility of cell division into substantially unequal parts. Asymmetry of distributions, in our opinion, is due to the fact that in the considered cellular structures, the minimum cell area is limited, not by zero, but by a specific positive value. We have also noticed that the maximum spread of areas strongly fluctuates and can noticeably differ in structures with similar morphology. Therefore, it is reasonable to characterize a cellular structure with the average area *S*_*av*_ of its cells and the average spread *ΔS*. By definition, *ΔS* is the difference between smallest and largest cell areas among the half of the cells with the areas closest to the *S*_*av*_ value. We then introduce a dimensionless effective spread Δ*S*/*S*_*av*_.

Structural data on the investigated epithelia are presented in Table I. The data in the last two lines are averaged for all the, respectively, synchronized (**<Hela**_**syn**_**>**) and non-synchronized (**<HeLa>**) HeLa samples studied. Note, that in 2-4^th^ columns, the values are weighted arithmetic means. Namely, the *S*_*av*_, Δ*S*/*S*_*av*_ and *P*_*6*_ values are weighted by the numbers of *N*_*vor*_ in each line.

The average cell area in the HeLa non-synchronized epithelium is larger than in the synchronized one (see Table I) since most of the cells in the latter belong to the second generation, with similar time elapsed after the division process. In all the examined specimens of both types, the distribution of cell areas was close to Gauss type. In HeLa synchronized epithelium, the spread in the average cell areas is apparently due to the growth of samples upon coverslips with an uneven surface. Therefore, the cellular monolayer undergoes a strain, which we consider to be homogeneous in the image size scale. The spread in the average cell areas between HeLa samples can be also related to another experimental feature. To prevent the formation of multilayers, the growth is stopped just before the total confluence of the cells. As a result, small and relatively sparse empty areas appear (see Methods). Note also that the averaged value < Δ*S*/*S*_*av*_ > is 7% larger in the HeLa non-synchronized data set. However, the averaged DOPTs remain very close, with the differences in probabilities Δ*P*_*i*_ ≲ 0.01 (see Fig. 1g).

### Model of random polygon packing

As we have already mentioned, synchronized cell monolayers are beyond the scope of applicability of the theory^**11**^. Inapplicability of this theory for the non-synchronized case is evidenced by our finding that DOPTs in both types of monolayers are very similar. Differences in structural parameters of different samples, the Gaussian nature of the cell area distributions, and their specific asymmetry lead to the hypothesis that in the hyperproliferative epithelia the cell packings are close to random but, nevertheless, satisfy the geometric constraint associated with the existence of minimal cell size. Below we develop the theory of random polygonal packings and then test our hypothesis on the random nature of the epithelial cancer cell monolayers.

Let us consider the random distributions of points with the minimal allowed distance between them, *d*_*min*_. The *Δ* value (Eq. 1) is exactly equal to 0 for any periodic arrangement of particles. Therefore, we will construct a model structure from *N* polygons in the square fundamental region that is repeated periodically. At the beginning, *N* points are randomly and sequentially inserted into the region with area *A*_*r*_. If the distance between the point that is currently being placed and any of the points placed earlier is less than the distance *d*_min_, then this point is deleted, and the process of random insertion is repeated. Next, after the placing of all *N* points, the Voronoi tiling is constructed with the nodes at these points.

The densest (fully ordered hexagonal) packing of particles corresponds to the case when *d*_*min*_= *d*_*hex*_, where 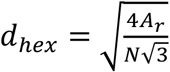, both *A*_*r*_ and *N* tend to infinity. It is almost impossible to obtain the strongly ordered structures by random placement of particles. For example, if *N* ∼ 5000, then about 5×10^5^ attempts to place points are necessary to generate a packing with *d*_min_∼0.758 *d*_hex,_ and containing ∼50% of hexagons. Figure 2a-f shows examples and area distribution histograms of random packings with different ratios *η*=*d*_min_/*d*_hex_ and different degrees of hexagonality. The plots in Fig. 2g are calculated up to *η*=0.75, since at larger *η* the calculation time increases sharply. Note also, that even at constant *η*, the algorithm generates packings with slightly different morphological properties. However, when *N* ∼ 5000, in the region of *η*>0.4, the ratio Δ*S*/*S*_*av*_ and probabilities *P*_*i*_ are reproduced with standard deviations smaller than 0.01 (see Fig. 2g).

**Fig. 2.**
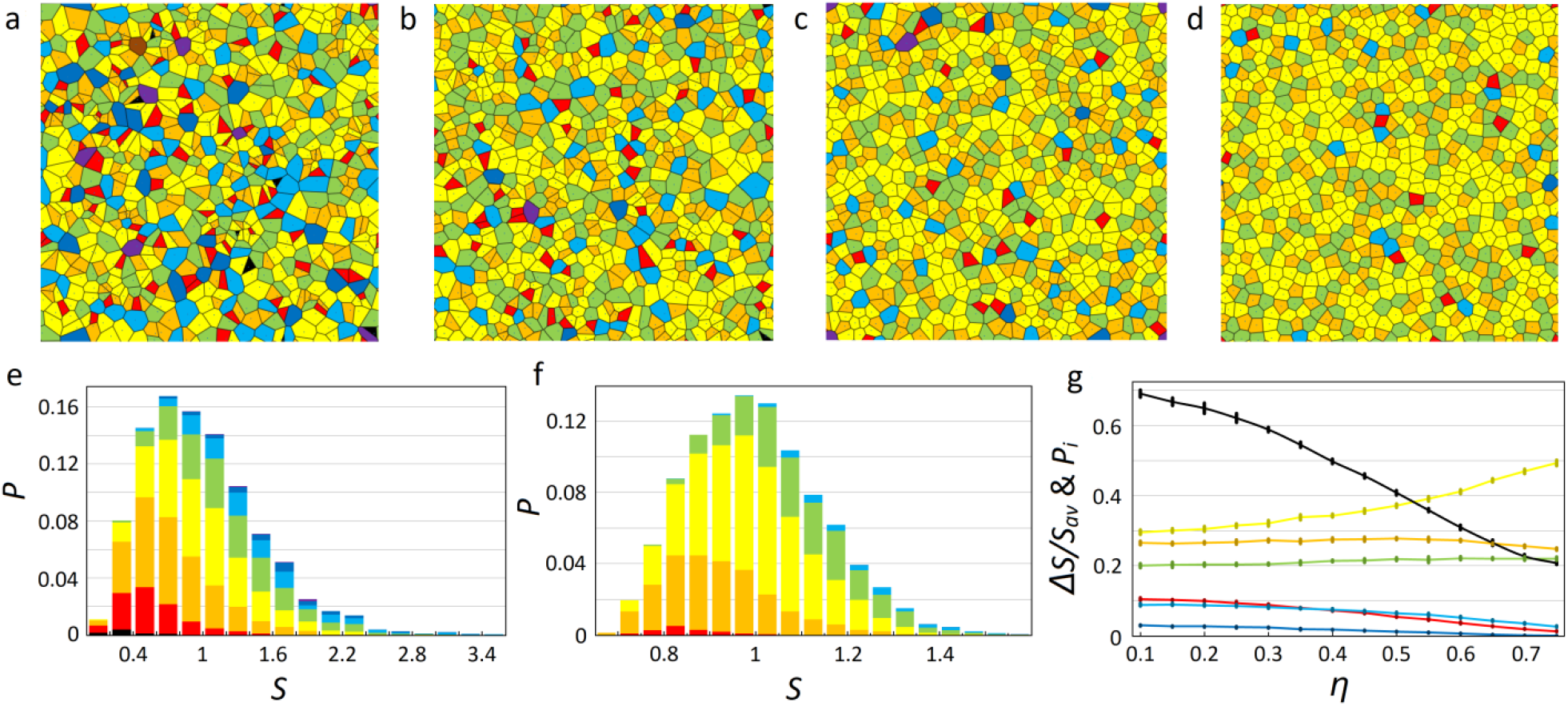
Examples of random packings and their geometric characteristics. **a-d** Packings obtained at *η* values equal to 0.1, 0.4, 0.6, and 0.758, respectively. It can be observed how the fraction of hexagonal cells grows with increasing *η* (these cells are shown in yellow; the coloring is the same as in Figs. 1c-f). **e-f** Area distribution histograms obtained for *η* = 0.1 and 0.758. For both cases, the value of *S*_*av*_ is renormalized to 1. **g** Dependences Δ*S*/*S*_*av*_ and *P*_*i*_ on *η*. Plots of Δ*S*/*S*_*av*_, *P*_*6*_, *P*_*5*_, *P*_*7*_, *P*_*4*_, *P*_*8*_, *P*_*9*_, calculated with the step *η*=0.05, are colored with black, yellow, orange, green, red, light blue and dark blue colors, respectively. Probabilities *P*_*3*_, *P*_*10*_ and *P*_*11*_ are too small to be shown in the chosen scale. Each center of vertical bars represents the averaging of 10 calculations at *N* = 5000. The bar sizes denote the standard deviations obtained for the calculations.

The proposed random packing model is in perfect agreement with the averaged structural data for the non-synchronized and synchronized HeLa epithelial cell monolayers (see the last two lines of Table I (**<Hela>** and **<Hela**_**syn**_**>)** and the histograms in Fig. 1g). Indeed, as presented in Fig. 2g, the change in Δ*S*/*S*_*av*_ from 0.4 to 0.43 corresponds to the variation of *η* from 0.5 to 0.47, respectively. However, in this region, the slope of the plot *P*_*6*_(*η*) is small, and the values of *η* correspond to very close DOPTs with *P*_6_ ≈ 0.35 ÷ 0.37. The deviations of other averaged probabilities (see Fig. 1g) from their theoretical values also do not exceed 0.02, which is within the spread of model calculations at *N* ∼ 5000.

The random packing model can also explain the scatter Δ*S*/*S*_*av*_ of and *P*_*6*_ values between different samples presented in Table I. Note that these monolayers contain, on average, slightly less than 500 cells, and such a small value of *N* increases the morphological inhomogeneity of the generated packings. To justify this, one can perform a series of computations with the appropriate inputs: the number of calculations should not be less than the total number of monolayers, and *N* is equal to *N*_*vor*_ in the sample. As a result, it is highly probable that a packing that deviates from the mean by about (or even more) than the considered sample will be generated.

As we already mentioned, small and relatively sparse empty domains are present in non-synchronized HeLa monolayers. We decided to evaluate the impact of these domains on the obtained results. For this purpose, on the assembled images, we selected 40 smaller regions without empty areas. These rectangular regions, with the perfect structure, contained from 36 to 76 cells. When averaging over these regions, the values of *S*_*av*_ were found separately, and cells for which it was impossible to determine the number of nearest neighbors were not considered. This treatment resulted in Δ*S*/*S*_*av*_ ≈ 0.394 and *P*_6_ ≈ 0.38. Comparing these values with those for averaged **<HeLa>** and using the graphs shown in Fig. 2g, we observe that the decrease in Δ*S*/*S*_*av*_ corresponds to the small increase in *P*_6_, and the order is still properly described by the developed theory of random polygonal packings.

Thus, we conclude that both types of considered hyperproliferative epithelia represent the random packings, while the main structural difference between the non-synchronized and synchronized HeLa monolayers consists in different values of *S*_*av*_.

## Discussion

It is important to discuss how a healthy proliferative epithelium distinguishes from random cancer monolayers and to propose a physical mechanism underlying this difference. As one can see from Fig. 2g, at *η* ∼ 0.62÷0.7, the model of random polygon packing reproduces perfectly the DOPTs^**8-11**^ typical of many plant and animal proliferative epithelia: *P*_4_ ≈ 0.02 ÷ 0.03, *P*_5_ ≈ 0.25 ÷ 0.27, *P*_6_ ≈ 0.41 ÷ 0.47, *P*_7_ ≈ 0.22, *P*_8_ ≈ 0.03 ÷ 0.05, and *P*_9_ ≈ 0.001 ÷ 0.006. A more detailed analysis, however, shows that the proposed approach leads to a value of Δ*S*/*S*_*av*_ that is slightly smaller than the one observed experimentally. In particular, after analyzing the images of the *Cucumis* proliferative epithelium^**8**^, we have estimated the value of Δ*S*/*S*_*av*_ as 0.29, which in the random polygon packing model corresponds to *η* ∼ 0.63 and *P*_*6*_ ≈ 0.43 instead of the observed value *P*_*6*_ ≈ 0.47^**8**^.

Let us consider the situation in more detail using our experimental data on healthy proliferative Human Cervical Epithelial Cells (HCerEpiC). Figure 3 (a-b) demonstrates a typical sample of this epithelium with the corresponding Voronoi tessellation. Forty analogous experimental images previously obtained in our laboratory represent relatively small separate fragments showing simultaneously from 30 to 54 cells. Processing of these images, as described above, leads to the following structural data: *S*_*av*_ ≈3.0×10^3^ μm^2^, Δ*S*/*S*_*av*_ ≈ 0.43, *P*_4_≈0.04, *P*_5_≈0.30, *P*_6_≈0.41, *P*_7_≈0.20, *P*_8_≈0.04, and *P*_9_≈0.01. It is interesting to note that the deviations of these *P*_i_ values compared to those obtained for the *Xenopus* frog^**11**^ are less than 2%.

**Fig. 3.**
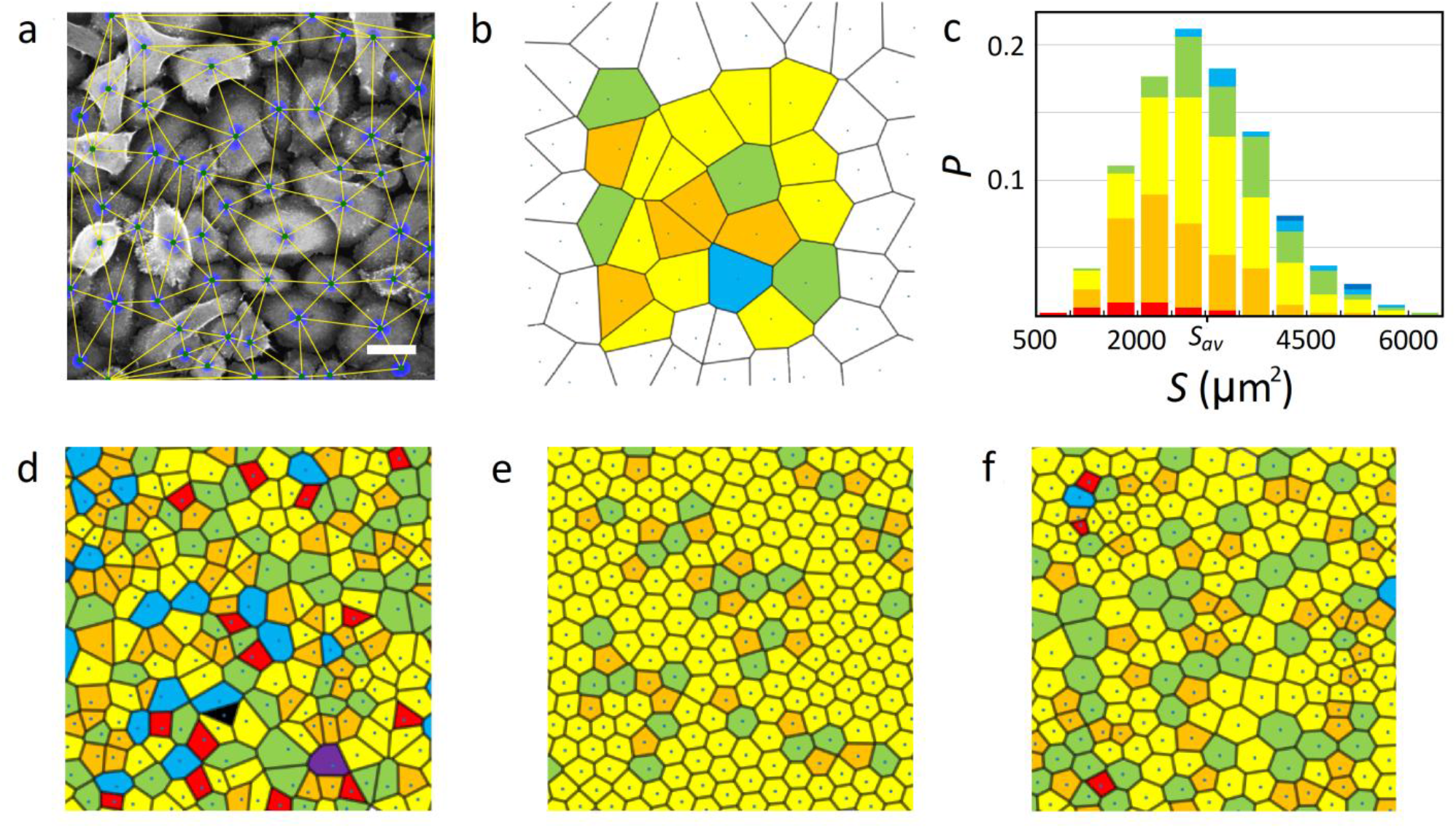
Structural characterization of HCerEpiC epithelium. **a** Typical sample of the epithelium with the superimposed triangulation; white scale bar is 50 μm. **b** Voronoi tessellation for the sample. A cell was taken into account in the statistical analysis if its shape could be determined unambiguously and the cell vertices were not too close to the border of the image (see Methods). Such cells are colored. **c** Averaged histogram of cell areas in forty samples; **d** Voronoi tiling for a random polygonal structure with the same Δ*S*/*S*_*av*_ value and very similar to (**c**) histogram of the cell area distribution. Polygons with 3-11 sides are colored with black, red, orange, yellow, green, light blue, dark blue, purple and brown colors, respectively. **e-f** More ordered structures obtained from **d** by minimizing the energy of elastic intercellular interaction (see main text). In (**e**) Δ*S*/*S*_*av*_ ≈ 0.14 and *P*_6_ ≈ 0.8, while for (**f**) *P*_6_ ≈ 0.55 and the value of Δ*S*/*S*_*av*_ is approximately equal to that in the HCerEpiC epithelium and random packing (**d**).

In the model of random packing, see Fig. 2g, the value Δ*S*/*S*_*av*_ ≈ 0.43 corresponds to *P*_6 ≈_0.36 instead of *P*_6 ≈_0.41 observed in HCerEpiC. Also, we have considered the fact that, due to the influence of the substrate on which the HCerEpiC monolayers were grown, the *S*_*av*_ value can be different in different fragments. With the averaging method taking this difference into account, the ratio Δ*S*/*S*_*av*_ decreases by ∼17%, corresponding to P_6_≈0.37, nevertheless, this slightly increased probability is still smaller than the observed value. Consequently, based on the *Cucumis* and HCerEpiC examples, we can conclude that in normal proliferative epithelia the value is greater than the one predicted by the random packing model for the same Δ*S*/*S*_*av*_ ratio.

Note that the mitosis rate in HeLa cells is ∼5.5 times higher than in HCerEpiC cells, while the rate of apoptosis is the same in both cases (See Supplementary Fig. 1). Due to this difference, for the same number of seeded cells, the monolayers are at confluence after 2 days for HeLa cells compared to 4 days for HCerEpiC ones. Thus, we can assume that the *P*_6_ increase in healthy epithelium is associated with the lower rate of cell division, however, intercellular interactions are also very important for this phenomenon.

To justify the latter statement, we recall that ordering of equivalent particles, retained on a planar surface and interacting with each other by very different pair potentials, readily leads to the formation of a simple hexagonal order^**30-32**^. In our opinion, the mechanism of the *P*_6_ increase in healthy proliferative epithelia is similar and caused by minimization of the elastic energy associated with the mechanical interaction between cells. Namely, appearance and non-uniform growth of new cells result in internal local stresses, which can relax due to cell motility, increasing, in average, the hexagonal coordination.

In the literature, we can find many different biophysical approaches (see some examples^**3,7,33-42**^) describing the growth, mitosis, motility, and interactions between cells. However, as far as we know, correlations between the cell area distribution and DOPT have never been investigated. Even cell area distributions in the numerous discussed epithelia are poorly studied. While undisputedly representing an important area for future researches, collection of new experimental data, their analysis and subsequent development of known or novel microscopic theories lay beyond the scope of this work. Below, we consider only a very simple model demonstrating that mechanical interactions between cells can order the epithelial structure by increasing the number of cells with six neighbors.

The DOPT observed in the epithelium HCerEpiC corresponds to the random polygonal packing with *η* ≈0.62, while the observed Δ*S*/*S*_*av*_ matches *η* ≈0.53. In the frame of the random packing model let us generate a Voronoi tessellation with *η* ≈0.53 (see Fig. 3d). This random structure yields a distribution of cell areas very close to that of the HCerEpiC epithelium (see Fig. 3c). Let us consider the nodes (shown with black points) of the tessellation as the centers of cells and introduce the intercellular interaction using the usual Lennard-Jones energy^**43**^:

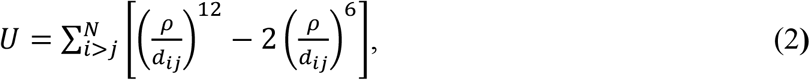

where *ρ* is the distance at which two-cells interaction reaches its minimum, *d*_*i,j*_ is the distance between the *i*-th and *j*-th nodes of Voronoi tessellation, *N* is the number of nodes. This energy has previously been used to model the cell-to-cell interactions in different studies^**28,44,45**^. Recall that the first term in Eq. (2) describes the contact repulsion, while the second one is responsible for the particle attraction at long distances. Since the cells can be of different size, we replace *ρ* with the sum of effective radii *r*_*i*_ and *r*_*j*_ : *ρ= r*_*i*_ + *r*_*j*_ Switching on the interactions described by energy (2) corresponds to the appearance of local stresses, while the minimization of energy (2) leads to their relaxation inducing some motion of the nodes.

Before minimizing the energy (2), we need to express the effective radii *r*_*i*_ in terms of the parameters characterizing the considered polygonal tiling. We assume that the total area of the structure after the energy minimization should be close to the initial one. By relating the effective cell radii *r*_*i*_ to the Voronoi cell areas *S*_*i*_ as 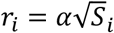 and minimizing (2) by the standard gradient descent method, we obtain such ordered structure (see Fig. 3e) with the value Δ*S*/*S*_*av*_ ≈ 0.14 and P_6_ ≈ 0.8 provided *α* ≈ 0.5 (for the packing of regular hexagons 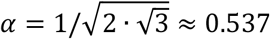). Note that the energy minimization decreases the initial Δ*S*/*S*_*av*_ value significantly. In order to conserve the halfwidth we express *r*_*i*_ in a more complicated phenomenological way as 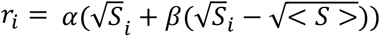, where < *S* > is the average area of the Voronoi cells. Then after the energy minimization both the total area and the initial value of Δ*S*/*S*_*av*_ ≈ 0.43 are conserved provided *β*≈2.2. Let us stress that the structures (see Fig. 3e, f) obtained by energy minimization are essentially more ordered than the epithelium of HCerEpiC as for the first structure *P*_*6*_ ≈ 0.80, and for the second *P*_*6*_ ≈ 0.55. Thus, mechanical interaction between cells, together with the cell motility, can significantly increase the proportion of 6-valent cells. Thus, the stress relaxation makes the normal proliferative epithelium more ordered in comparison to the almost random packings of cancer cell monolayers, where the rapid increase in cell number and the rapid growth of cell sizes prevent the minimization of free energy, and the cells tend to form random structures.

In contrast to the tumoral monolayers, in normal epithelia, the mitosis does not increase the disorder substantially and only prevents the growth of *P*_*6*_ up to the values estimated from the energy minimization. Therefore, the relation between the mitosis rate and the rate of epithelial ordering due to intercellular interactions controls the fraction of 6-valent cells, which, in turn, mainly determines all other *P*_*i*_ values. In this context, it should be noted that the epithelia with *P*_6_ ≈ 0.6 ÷ 0.9 are also observed in nature. For example, after the mitosis in the wings of a *Drosophila* stops (and before the start of hair growth), the equilibrium changes and a sharp increase in the proportion of 6-valent cells occurs^**46**^. In our opinion, this process is mainly associated with energy minimization and relaxation of accumulated local mechanical stresses. Thus, the identified mechanism, which works well in normal epithelium, is similar in many respects to what occurs in metals and alloys upon annealing, when the structure gets rid of defects due to interatomic interactions and diffusion of atoms.

In conclusion, for the first time, we have demonstrated and detailed the significant difference in topology of the cancerous and normal epithelial monolayers. In epithelial cancer monolayers, the cell arrangement is close to random but, nevertheless, satisfies the geometric constraint associated with the existence of minimal cell size. This type of the cell order is controlled with a good accuracy by only one dimensionless parameter, namely the normalized halfwidth Δ*S*/*S*_*av*_ of the cell area distribution. This happens because the cancer cells divide so quickly that the relaxation processes associated with the minimization of free energy and objective to increase the number of 6-valent cells do not have time to order the monolayer. As a consequence, the growing disorder predominates over the intercellular interactions and cell motility. On the contrary, in normal proliferative and non-proliferative epithelia, the tendency to order is substantial or even decisive. Our study of the tumor models can be very useful for the biophysical and biomedical research communities and may provide novel tools to test anti-cancer drugs.

## Methods

### Cell line growth and synchronization procedure conditions

Human cervical cancer cell line HeLa was purchased from the American Type Culture Collection (ATCC; Manassas, VA, USA) and maintained in DMEM, high glucose (Dulbecco’s Modified Eagle Medium) containing 5% heat-inactivated foetal bovine serum and supplemented with GlutaMAX™ (Gibco Life Technologies), penicillin (100 units/mL), and streptomycin (100 µg/mL). Normal primary cervical epithelial cells (HCerEpiC) isolated from human uterus were purchased from ScienCell Research Laboratories (Clinisciences S.A.S., Nanterre, France). HCerEpic cells were grown in Cervical Epithelial Cell Medium according to the manufacturer’s instructions. Cells were incubated in a humidified incubator at 37°C in 5% CO_2_.

For confocal microscopy analysis of non-synchronized cells, HeLa or HCerEpiC cells were seeded on glass coverslips (12mm diameter round) coated with 10µg/ml of poly-L-Lysine (P4707, Sigma) at 7·10^4^ cells/coverslip in a 24-well culture plate. Confluency of the monolayer was achieved 48 h and 4 days later for HeLa and HCerEpiC cells, respectively. HeLa cells were synchronized in G0/G1 phase of the cell cycle using the double thymidine block procedure. The day before the first thymidine block, cells were seeded on poly-L-Lysine treated glass coverslip at a density of 7.5×10^4^ cells/ coverslip. The next day, 2.5 mM thymidine was added for 16 h (first block). Then cells were washed with phosphate-buffered saline (PBS) and incubated during 8 h without thymidine. Lastly, the second thymidine block was applied for 16h. At the end of the second thymidine block, cells were washed and incubated for 2 additional hours, after which cells were treated with antibodies and analysed by confocal microscopy. Following this procedure, cells are fully confluent and cell synchronisation in G0/G1 is about 92%. This was confirmed by cytofluorometry analysis (FACS) (see Supplementary Fig. 2).

### Immunocytochemistry and fluorescence microscopy

Cells cultured on glass coverslips were fixed with 4% paraformaldehyde in PBS for 20 min, and washed in Tris-buffered saline (25mM Tris pH7.4, 150mM NaCl) (TS) for 10 min. After permeabilization with 0.2% Triton X-100 in TS for 4 min, non-specific binding was blocked with 0.2% gelatin from cold water fish skin (#G7765 Sigma-Aldrich Chimie, Lyon) in TS for 30min. Cells were incubated with primary antibodies in blocking buffer for 1h and then washed 3 times with 0.008% TritonX-100 in TS for 10 minutes. Rabbit anti-ezrin antibody^**47**^ was used to visualize cell body and membrane. Cells were incubated for 30 minutes with Alexa-Fluor 488 - labelled secondary antibodies (P36934-Molecular Probes, InVitrogen) in blocking buffer. After rinsing in washing buffer, cell nuclei were stained with 1 µg/ml Hoechst 33342 (62249-Thermo Scientific Pierce) in TS for 5 minutes. Finally, coverslips were mounted with Prolong™ Gold Antifade (P36934-Molecular Probes, InVitrogen) and examined under a Leica TCS SPE confocal microscope equipped with a 25X/0.75 PL FLUOTAR oil objective (HCerEpiC) and a 40X/1.15 ACS APO oil objective (HeLa) (Fig. 3 and Supplementary Fig. 1). In the case of HeLa cells (Fig. 1a) and synchronized HeLa cells (Fig. 1b), the analysis was performed with a Zeiss LSM880 FastAiryScan confocal microscope equipped with a 40X/1.4 Oil Plan-apochromat DIC objective.

Between 7 to 9 ‘z-stacks’ (0.457µm thickness each) were acquired per field and a 2D image was generated by applying Maximum Intensity Projection processing. For each coverslip, the acquisition pattern was 6 neighbouring images per row for a total of 2 or 3 rows. The resulting images (12 or 18 images) were adjusted for brightness, contrast and color balance by using ImageJ and assembled side by side in PowerPoint to reconstruct a cell monolayer consisting of N>500 cells.

### Image analysis

After determining the geometric centers of the cell nuclei, triangulation was performed by the Delaunay method^**48**^. Next, Voronoi tiling (which is dual to the Delaunay triangulation) was constructed and the areas of the epithelial cells were calculated as the areas of Voronoi cells. Obviously, for a correct statistical analysis, it is necessary to discard the cells (located too close to the image border), the number of neighbors for which cannot be determined. Note that even if it is possible to construct a closed Voronoi cell, then it is also necessary to check whether the cell polygon boundary can be changed by additional hypothetical nuclei lying directly outside the image border. Therefore, the center of a reliably constructed Voronoi cell should be located at least twice as far from the image border as any of the vertices of this cell. However, this method leads to the appearance of an excessive total positive topological charge, which is localized at the image border. On one hand, 4- and 5-valent cells have a smaller area^**8**^, while on the other hand, the smaller the cell located near the image border, the more chances its nucleus has to satisfy the selection criterion formulated above. This fact, when processing images with a small number of nuclei (about 40), leads on average to the formation of a 5% preponderance of the total positive topological charge (which is carried by 4- and 5-valent cells) over the total negative topological charge. This, in turn, leads to errors when constructing DOPTs diagrams and determining the value of *P*_6_ and Δ*S*/*S*_*av*_ To avoid preferential selection of small cells, we used additional cutting of the image borders. In the statistical analysis, we took into account only cells whose nuclei centers fall within the rectangle, which has maximum possible size and does not contain any nucleus with an uncertain coordination. Thus, it is possible to significantly reduce the total topological charge of the images and, accordingly, the error in the values of Δ*S*/*S*_*av*_.

## Data availability

Data supporting the findings of this manuscript are available from the corresponding authors upon reasonable request.

## Acknowledgments

M. M., V.M., and S.B. acknowledge the imaging facility MRI, member of the national infrastructure France-BioImaging infrastructure supported by the French National Research Agency (ANR-10-INBS-04, «Investments for the future»). S.R. and D.R. acknowledge the financial support from the Russian Foundation for Basic Research (Grant № 19–32–90134). We thank A Parmeggiani and I. Golushko for useful discussions.

## Author contributions

Several years ago, S.B. proposed to S.R. to think about the topological universality of epithelia. When S.R. proposed the model of random polygonal packings, S.B. initialized this study. M. M. performed the biological experiments and the confocal microscopy acquisition as well as assembly of images. K.F. realized the initial analysis of images. D.R. did most of the calculations and analysis. S.R. wrote with the essential help of S.B. and M.M. the main part of the text. V.M. proofread and corrected the article extensively. All authors participated in discussion and writing of the article.

## Additional information

Supplementary information is available for this paper at ***

