## Supplementary Fig. for "Cancer cell hyper-proliferation disrupts the topological invariance of epithelial monolayers"

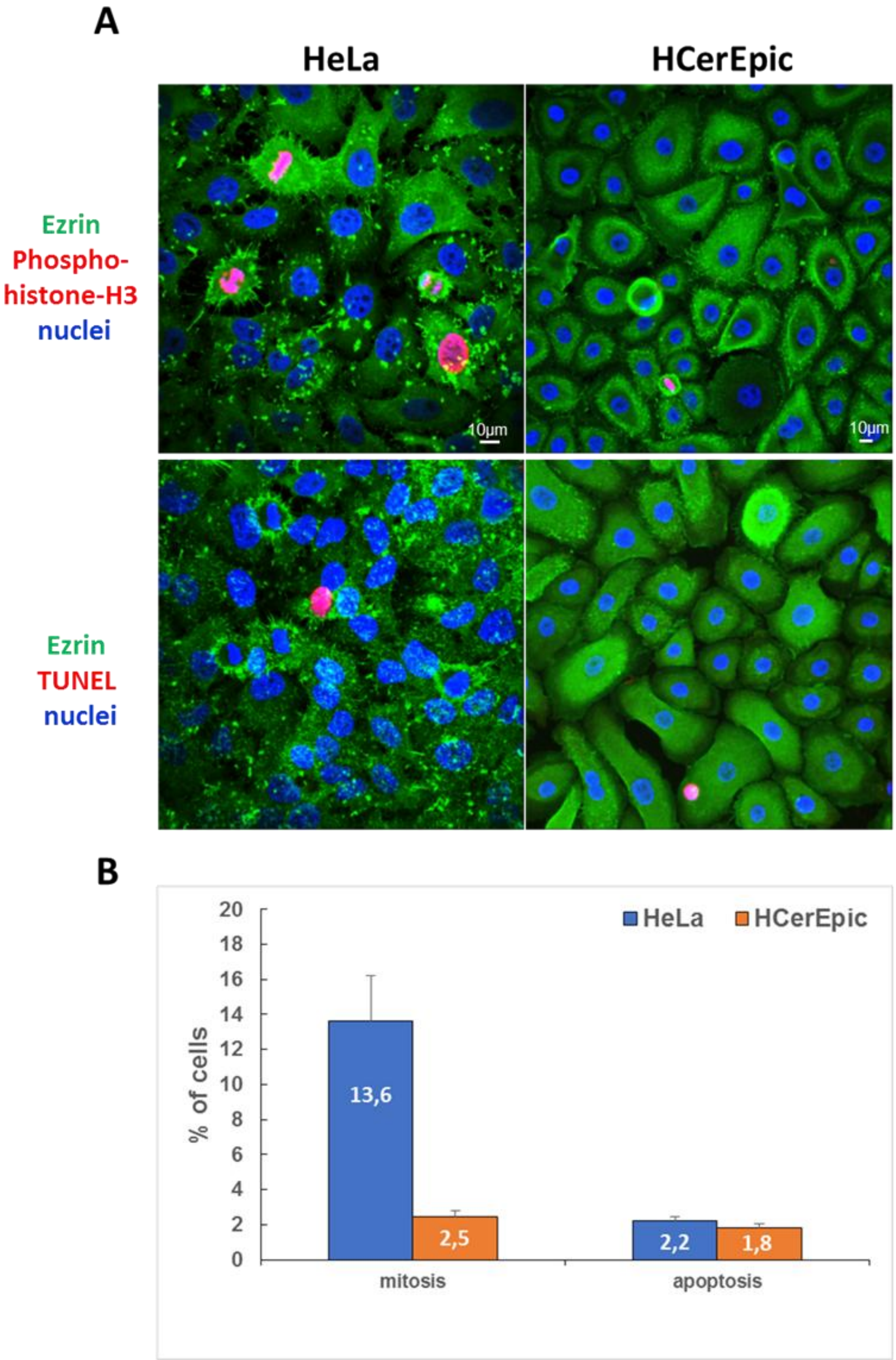


**Fig. S1. Percentage of HeLa and HCerEpic cells in mitosis and apoptosis.** Cells were grown on glass coverslips until confluency. After formaldehyde fixation, they were permeabilized and treated for further analysis by immunofluorescence confocal microscopy. Panel A shows representative images of cells in mitosis or apoptosis (pink labelling). Nuclei were stained by Hoechst (blue labelling) and an antibody against ezrin (green) was used for cells labeling. To determine the level of mitosis, Hela and HCerEpic were labelled with an anti-phosphohistone H3 (pSer10) antibody as a proliferation marker. This marker allows to visualize only the four actual phases of mitosis and late G2. Quantification of mitosis was obtained from 3 coverslips with 10 acquisitions per coverslip (HeLa: 2567 cells; hCerEpic: 3804 cells). Detection of apoptotic cells was obtained by labelling the cells with the In Situ Cell Death Detection Kit, TMR red (TUNEL)

according to the manufacturer. This kit is based on the detection of single- and double-stranded DNA breaks that occur at the early stages of apoptosis. For apoptosis, 10 images per coverslip were collected for a total of 10 coverslips analysed (HeLa: 874 cells; hCerEpic: 1345 cells). Histogram in B shows the percentage of cells in mitosis and apoptosis. HeLa cells are significantly more proliferative than HCerEpic cells (5.5 times more) whereas both cell lines have similar apoptotic level. Error bars are Standard Error of the Mean (S.E.M.)

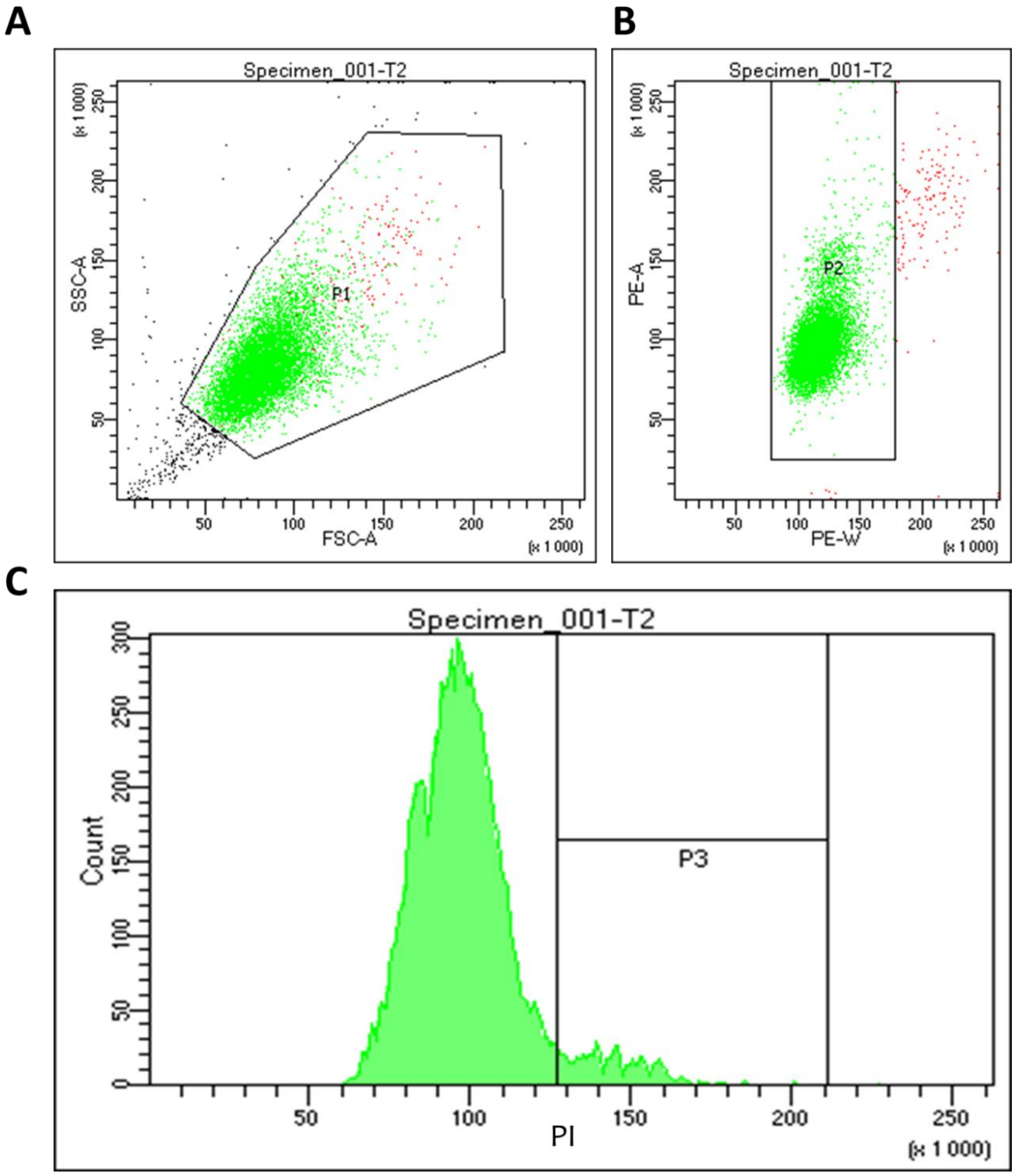


Tube: T2

| Population | #Events | %Parent | %Total |
|---|---|---|---|
| All Events | 10 000 | #### | 100,0 |
| P1 | 9 673 | 96,7 | 96,7 |
| P2 | 9 525 | 98,5 | 95,2 |
| P3 | 682 | 7,2 | 6,8 |

**Fig. S2. Flow cytometric analysis of synchronized HeLa cells.** HeLa cells were synchronized by a double thymidine block. After elimination of thymidine, cells were incubated 2 hours in complete medium. Supernatant containing apoptotic cells was discarded and adherent cells were detached by trypsin. Cells were washed once with PBS. After centrifugation, the pellet was resuspended at $1.10^6$ cells in 500 µl of PBS and fixed with cold ethanol at a final concentration of 70% during 2 h at 4°C. After one wash with PBS, cellular DNA was stained by incubation in PBS containing 100 µg/ml RNase DNAse-free and 40 µg/ml Propidium Iodide (PI). Plot in A allows exclusion of cells debris and selection of P1 population (P1 96.7%). Pulse shape analysis in B is used to exclude clumps and cell doublets from the analysis of P1 population (P2 population 95.2%). Plot in C shows the resulting PI histogram taking into account results from plots A and B. P3 population (6.8% of P2 population) is representative of cells in S phase and G2/M. Therefore, ~92% of HeLa cells are synchronized in GO/G1 after the double thymidine block.
